# Spatial distribution of *Salmonella enterica* in poultry shed environments observed by intensive longitudinal environmental sampling

**DOI:** 10.1101/547539

**Authors:** Helen K Crabb, Joanne Lee Allen, Joanne Maree Devlin, Colin Reginald Wilks, James Rudkin Gilkerson

## Abstract

During a series of epidemiological studies to investigate the transmission of salmonellae within poultry environments, intensive longitudinal sampling within caged sheds revealed that both the number and location of sample collection within this environment were important to ensure the best chance of detecting *Salmonella* spp. Multiple serovars of *Salmonella enterica* subspecies *enterica* were detected in each shed; 5% of all samples contained more than one serovar. Samples collected on the north side of the shed (OR = 1.77, 95% CI [1.17, 2.68]), on the sheltered side of the shed (OR = 1.90, 95% CI [1.26, 2.89]) and during winter (OR = 48.41, 95%CI [23.56, 104.19]) were more likely to be positive for salmonellae. The distribution of salmonellae within a shed was not homogenous, as differences were identified in the within-shed distribution of *Salmonella enterica* subspecies *enterica* serovar Typhimurium (*χ*^2^ (27, 1,538) = 54.4, P < 0.001), and *Salmonella enterica* subspecies *enterica* serovar Infantis (*χ*^2^ (27, 1,538) = 79.8, P < 0.0001). The difference in sample prevalence by serovar and spatial location within a shed indicates that there are important shed micro-environmental factors that influence survival and/or distribution of salmonellae. These factors should be taken into consideration when undertaking environmental surveillance for salmonellae in flocks housed in caged sheds.

**IMPORTANCE:** Routine epidemiological surveillance for salmonellae in poultry relies initially on environmental sampling. Our study confirmed that the sampling methodology conducted within a poultry environment is a non-trivial part of sampling design. The spatial locations of sample collection within a shed revealed differences in both the sample prevalence and serovar of salmonellae detected.

## INTRODUCTION

The routine surveillance for salmonellae in poultry typically involves environmental sampling because it has long been established as both the most cost effective and sensitive method for detection (1) when compared to individual bird sampling (2–4). Environmental sampling is indicative of flock infection with salmonellae, while the level of environmental contamination (both semi-quantitative and sample prevalence) and egg prevalence may be correlated with prevalence of infection in the flock (5, 6). As flock prevalence declines the number of environmental samples required to detect infection must increase to maintain adequate test sensitivity (7, 8). Importantly, animal welfare and production effects associated with live bird handling are minimised, allowing environmental sampling to be conducted at frequent intervals with little to no flock interference.

A variety of environmental sampling methods and strategies have been developed for the detection of salmonellae in poultry environments (1, 9, 10) and the principal features of these methods have been incorporated into national surveillance and control programmes (11–15). While extensive knowledge has been gained regarding environmental sampling for the identification of infected flocks for control purposes, few studies have described intensive longitudinal sampling for salmonellae within infected flocks or environments (16–19). It is known that salmonellae will survive for extended periods of time in contaminated environments under optimal conditions. An increase in number of positive samples has been observed in winter periods (9), although it is not clear whether this is due to increased shedding from infected birds, prolonged survival during cooler weather or multiplication of salmonellae in the environment (18). In addition, multiple *Salmonella* spp. may be detected in flocks at a single environmental sampling event (19).

Despite extensive discussion regarding sample type and sample pooling, the method of sample site selection and sample collection within a cage shed environment is infrequently described or discussed (9, 20). The method of choosing environmental sites for sampling within a poultry cage shed environment is a non-trivial part of surveillance design. Important information about the distribution of salmonellae and other pathogens within the environment and how this may affect surveillance strategies beyond simple presence or absence of detection is largely unknown or undescribed in the literature. No studies have repeatedly sampled flocks within the same environment at a high frequency (3 weekly) for the lifespan of the flocks contained therein. Thus, it is not known what, if any, environmental or flock factors may influence the repeated detection of salmonellae during the life of the flock, or what influence post-cleaning decontamination may have on subsequent flock infection or the detection of environmental *Salmonella* spp.

In veterinary science, prevalence calculations typically use herd or flock size to calculate the appropriate sample size to detect infection at a given prevalence (21), so that the sampling strategy chosen is sufficiently powerful to provide sufficient confidence of detecting infection at this determined prevalence (22). When extrapolating this information to environmental sampling for the detection of a specific pathogen, thus indirectly sampling the flock, several unknowns are encountered. Does repeated detection reflect flock shedding of salmonellae, or rather the repeated detection of the same salmonellae persisting or multiplying in the contaminated environment? Is the failure to detect salmonellae due to inadequate sample site selection, insufficient sample size, or differences in distribution within a caged shed environment? How does one ensure that sufficient samples are collected that adequately represent both the environment and the distribution of infection within a flock, assuming infection may be heterogeneously distributed? Our investigations and subsequent findings, may be useful to others when designing sampling approaches for *Salmonella* spp. detection in caged poultry flocks.

## RESULTS

### Sample size estimation

The number of samples required to be 95% confident that the estimate of prevalence was within 5% of the true population prevalence for each unit of interest (1. Birds, 2. Cages or 3. Environment) are summarised in TABLE S1. At least 28 cages or environmental samples are required to estimate the *Salmonella* spp. sample prevalence with sufficient confidence at the lowest design prevalence. At the lowest design prevalence, at either diagnostic test sensitivity, the number of birds to be sampled was greater than the precision of the sample calculation and unable to be calculated.

### Surface area sampled

On each sampling occasion, between 208 – 224 m^2^ of surface area per shed was sampled. The total surface area of each shed sampled by sample type is described in TABLE S2.

### Sampling Results

A total of 19 caged flocks were sampled on 105 occasions, with 2,879 samples collected. Overall, 37% of all samples collected were positive for *S. enterica*, with more samples obtained from conventional cages testing positive compared to those collected from colony cages. *S.* Typhimurium and *S.* Infantis serovars were the most frequently detected serovars, comprising 9.8% and 14.9% of the isolates respectively. In the colony cages *S.* Typhimurium and S. Infantis were the only serovars detected but in the conventional cages other *S. enterica* serovars (including *S.* Singapore*, S.* Agona*, S.* Virchow) were also detected (TABLE 1)

### Predictive value of sampling method

The positive and negative predictive values of the combined sampling methodology for *S. enterica* were high (TABLE 2). Importantly, the negative predictive value of the combined sampling method was high regardless of the serovar. However, the positive predictive value of the sampling method differed between *S. enterica* serovars, with the probability of detecting S. Typhimurium lower than *S.* Infantis, reflecting the enrivonmental prevalence of the respective serovars.

### Sampling events

All flocks were sampled at least twice, the median number of sampling events for all flocks was 3 (range 2 - 15). Twelve of the 19 flocks were longitudinally sampled; 8 flocks at least monthly until 40 weeks of age, and 4 flocks (A, B, C, D) housed in colony cage sheds, at 3 weekly intervals until flocks were ∼65 weeks of age. All flocks were detected positive for *S. enterica* on at least one sampling occasion (TABLE 3) and multiple *S. enterica* serovars were detected in all sheds.

**TABLE 1.**
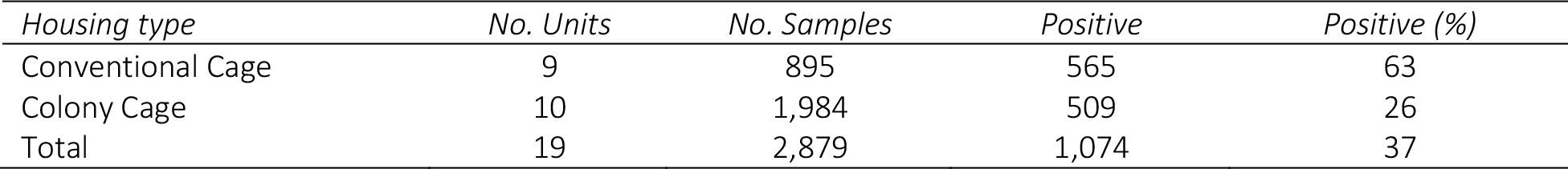
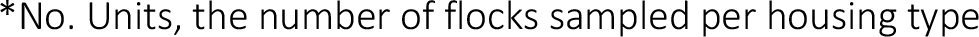
Summary of environmental testing results by housing type

| Housing type | No. Units | No. Samples | Positive | Positive (%) |
| --- | --- | --- | --- | --- |
| Conventional Cage | 9 | 895 | 565 | 63 |
| Colony Cage | 10 | 1,984 | 509 | 26 |
| Total | 19 | 2,879 | 1,074 | 37 |

\*No. Units, the number of flocks sampled per housing type

**TABLE 2.** Positive predictive values for the sampling methodology and the odds of detection for each sample type for *S. enterica*, *S.* Typhimurium and *S*. Infantis

|  |  | *S. enterica* | *S. Typhimurium* | *S. Infantis* |
| --- | --- | --- | --- | --- |
| Boot | OR | 7.69 (5.60 - 11.35) | 2.12 (1.36 - 3.65) | 5.14 (3.66 - 7.78) |
| Dust | OR | 3.84 (2.94 - 5.31) | 2.76 (1.91 - 4.32) | 2.98 (2.20 - 4.31) |
| Manure | OR | 5.19 (3.94 - 7.28) | 2.94 (2.03 - 4.62) | 3.53 (2.59 - 5.17) |
| Egg | OR | 1.00 | 1.00 | 1.00 |
| All | P(B+\|A+)<sup>1</sup> | 0.984 (0.979 - 0.989) | 0.907 (0.855 - 0.959) | 0.943 (0.902 - 0.985) |
|  | NPV<sup>2</sup> | 0.994 (0.979 - 1.00) | 0.999 (0.994 - 1.00) | 0.998 (0.991 - 1.00) |

<sup>1</sup>PPV - P(B+|A+) probability that the sample type is positive if all the samples are positive
<sup>2</sup>NPV - P(B-|A-) probability that the sample type is negative if all the samples are negative

**TABLE 3.** Mean true sample *S. enterica* prevalence summarized by housing type and sampling event for each serovar

| *Housing Type* | *Sampling Events* | *Negative Sampling Events* | *Mean True Prevalence (S.E.) Sample Event* |  |  |
| --- | --- | --- | --- | --- | --- |
|  |  |  | *All S. enterica* | *S. Typhimurium* | *S. Infantis* |
| Conventional Cage | 31 | 1 | 0.64 (0.09) | 0.03 (0.03) | 0.17 (0.03) |
| Colony Cage | 74 | 14 | 0.26 (0.05) | 0.13 (0.04) | 0.14 (0.04) |
| All | 105 | 15 | 0.37 (0.05) | 0.10 (0.03) | 0.15 (0.04) |

The odds of a samples being positive for *S. enterica* varied substantially by shed (OR = [Range 3.04 – 144.53]), indicating substantial shed effect with regards to *S. enterica* persistence or survival within an individual shed, either within the birds or within the shed environment (TABLE S3).

Negative sampling events were more likely to occur in colony cage houses than conventional cage houses (OR = 60.12; 95% CI [7.39, 48.11], *χ*^2^ = 34.24, P < 0.001), but *S*. Typhimurium was slightly more likely to be detected in colony cage samples (OR = 2.19; 95% CI [0.89, 5.39], *χ*^2^ = 2.99, P = 0.08). *S.* Infantis was more likely to be isolated from positive samples, regardless of shed type (OR = 1.76; 95% CI [1.49, 2.08], *χ*^2^ = 45.40, P < 0.001).

### Post Cleaning

All sheds were either wet or dry cleaned prior to repopulation. Twelve flocks were sampled post cleaning and salmonellae were detected in 9 of 12 sheds. *S.* Typhimurium and *S.* Infantis were detected in 3 and 7 of the cleaned sheds, respectively. There was no statistically significant difference between post cleaning and production sampling, when the sample prevalence of *S. enterica* was estimated for each sampling event.

A significant difference was observed when the method of cleaning was considered. In sheds that were wet washed the sample prevalence was lower immediately post cleaning (T (17) = 3.54, P < 0.001) and over all sampling events (T (17) = 6.11, P < 0.001). A wet washed shed was more likely to have *S. enterica* negative sampling events during the subsequent flock production period than sheds that were dry cleaned only (OR 1.44; 95% CI [0.43, 4.82], *χ*^2^ = 0.35, P = 0.55), however this difference was not statistically significant.

### Sample Type

Results for all sheds aggregated by sample type and *S. enterica* serovar are presented in TABLE 4. The estimated true sample prevalence for all *S. enterica* serovars was 39%, with *S.* Infantis (14%) more frequently detected than *S*. Typhimurium (9%). When taking into account both the shed type and sampling event, all sample types were significantly better than egg belt samples (F (3, 2589) = 42.84, P < 0.001) for detecting S. enterica regardless of the serovar. For the detection of *S.* Typhimurium there was no statistically significant difference between sample types (Dust = Manure Belt = Boot Swab), however boot swabs were better than all other samples for detecting *S*. Infantis (Boot Swab > Dust = Manure Belt) (F (3, 2589) = 21.7, P < 0.001).

**TABLE 4.**
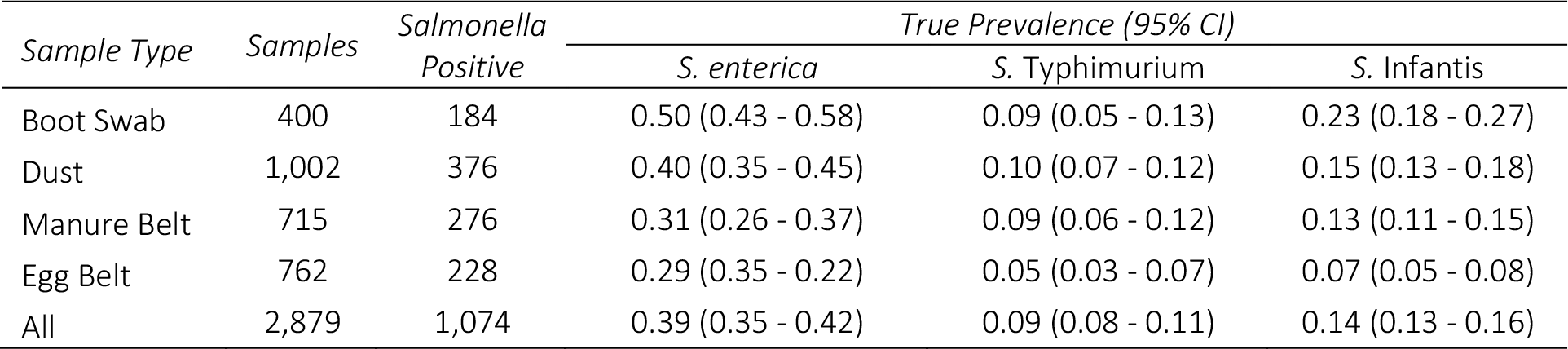
True sample *S. enterica* prevalence and 95% confidence intervals for all flocks and sample events by sample type for each serovar

### Other factors

Samples collected on the north side of a shed (P < 0.001), or on the aspect of a shed sheltered by another shed (between) (P < 0.001) were more likely to be positive for *S. enterica* (TABLE 5). There was a significant difference between flocks and sheds in the number of positive samples collected at each sampling event (P < 0.001), indicating that it is important to take into account the hierarchical effects of the sampling design.

**TABLE 5.** Summary of univariate analysis results of environmental variables associated with true sample *S. enterica* prevalence, aggregated for all sheds and flocks

| *Variable* | *Code* | *OR* | *95% CI* | $\chi^2$ | *DF* | *P-value* |
| --- | --- | --- | --- | --- | --- | --- |
| Flock | 1-19 | — | — | 117.1 | 18 | < 0.001 |
| Shed | 1-13<sup>1</sup> | — | — | 445.4 | 12 | < 0.001 |
|  | North | 1.33 | 1.12 – 1.57 | 10.7 | 1 | < 0.001 |
| Aspect | South | Reference | – |  |  |  |
| Betweenness | Yes | 1.96 | 1.64 – 2.33 | 56.9 | 1 | < 0.001 |
|  | No | Reference | – |  |  |  |
| Month | 1-12 | – | – | 196.7 | 11 | < 0.001 |
| Sampling Event | 1-15<sup>2</sup> | – | – | 200.4 | 14 | < 0.001 |
| Sample Type | Boot Swab | 2.42 | 1.85 – 3.17 |  |  |  |
|  | Dust | 1.63 | 1.31 – 2.03 | 45.4 | 3 | < 0.001 |
|  | Manure Belt | 1.71 | 1.35 – 2.15 |  |  |  |
|  | Egg Belt | Reference | – |  |  |  |
| Season | Autumn | Reference | – |  |  |  |
|  | Spring | 0.46 | 0.36 – 0.59 | 133.5 | 3 | < 0.001 |
|  | Summer | 0.74 | 0.59 – 0.94 |  |  |  |
|  | Winter | 2.02 | 1.57 – 2.59 |  |  |  |

<sup>1</sup>Table S3, <sup>2</sup>Table S4

When the month of sampling was considered, samples collected in June and August (P < 0.001) were more likely to be positive for *S. enterica*. There was a seasonal effect with samples collected in winter (P < 0.001) more likely to be positive than those collected in the other seasons (winter > spring > summer > autumn). Full results of the weather variables considered in the analysis are detailed in the supplementary material (Weather Results, TABLE S6, Table S7).

### Colony cage sheds

To further investigate the environmental factors affecting *S. enterica* prevalence and distribution in detail, the four most intensively sampled flocks were considered in depth. These flocks (A, B, C, D) were sampled on 13-15 occasions during the flock production period. A total of 1,538 environmental samples were collected; of these 24% were positive for *S. enterica*. Five percent of the samples contained more than one *S. enterica* serovar, but only *S.* Typhimurium and *S.* Infantis were identified in these sheds. In 3 of the 4 sheds *S.* Typhimurium was isolated more frequently (TABLE 6). The percentage of *S. enterica* positive samples (8 – 56%), varied by shed and serovar, and the difference in sample prevalence between sheds was statistically significant (*χ*^2^ (3, 1,538) = 246.8, P < 0.001).

**TABLE 6.**
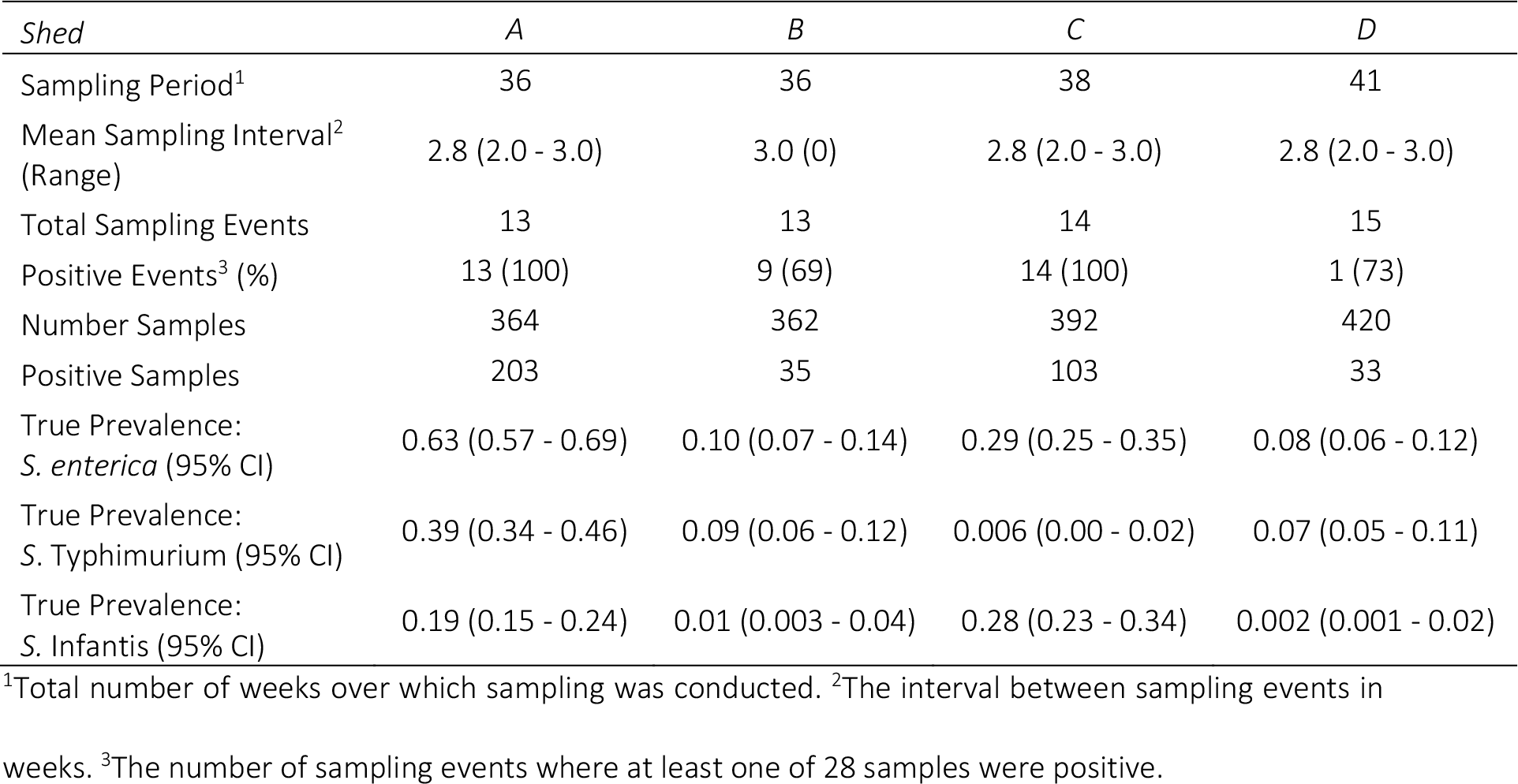
Summary of environmental sampling in the four intensively sampled colony sheds

All sheds were negative for *Salmonella* spp. prior to the onset of flock placement. *S. enterica* were detected in all sheds on most sampling events (69 – 100%). However, the number of sampling events that were positive for each sample type varied for any one shed (TABLE S4). The odds of a sample being positive for *S. enterica* varied by time (sampling event) with samples collected on the first sampling event significantly more likely to be positive than those collected later in the sampling period (TABLE S5).

Those sheds with fewer positive sampling events had fewer *S. enterica* positive samples. In sheds A and C (high environmental prevalence sheds) any combination of boot or dust sample would have detected that these were sheds positive for *S. enterica* on any sampling event. Whereas in sheds B or D (low environmental sample prevalence sheds) fewer than half the samples were positive on any sample event regardless of the sample type. Only by combining the results of each sample type on each sampling event did the probability of detecting *S. enterica* increase, 9– to 12– fold in sheds B or D respectively.

There was a significant difference between locations within a shed for the detection of *S. enterica* (*χ*^2^ (27, 1,538) = 96.6, P < 0.001). Samples taken on the north side of the shed (OR = 1.95, 95% CI [1.54-2.49]), and those collected from locations between sheds were more likely (OR = 1.92, 95% CI [1.52 - 2.45]) to be positive.

Thirteen of the 28 sampling locations were more likely to be positive for *S. enterica* regardless of the serovar (Table S6). *S*. Typhimurium was more likely to be found in 7 of the 28 locations (*χ*^2^ (27, 1,538) = 54.4, P < 0.001), and *S.* Infantis was more likely to be identified in 7 different locations (*χ*^2^ (27, 1,538) = 79.8, P < 0.001). The detection of a *S. enterica* positive sample at one location was not influenced by the detection of another positive sample in the same or similar spatial location (spatial autocorrelation, Moran I = −0.026, P = 0.343). The frequency that each serovar was identified at a location is illustrated in FIG. 3A, and the sample prevalence for each serovar, illustrated in a 2D space, is presented in FIG. 3B and FIG. 3C.

### Hierarchical mixed-effects multivariable modelling

The univariate results for all statistically significant explanatory variables considered for multivariable modelling are summarized in TABLE 7, a full description of all variables is available in TABLE S7. The final multivariable model considered the detection of a *S. enterica* positive sample, irrespective of serovar. Estimated regression coefficients for the final model and the variability of the random effects term are provided in TABLE 8.

**TABLE 7.**
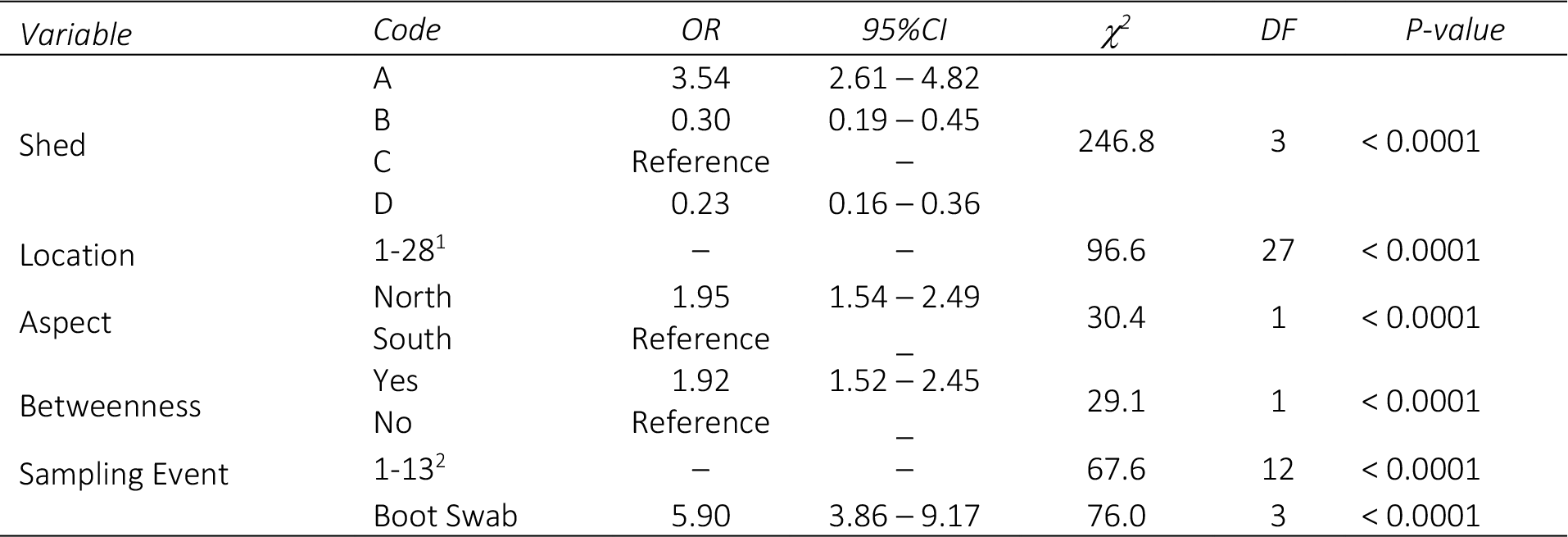
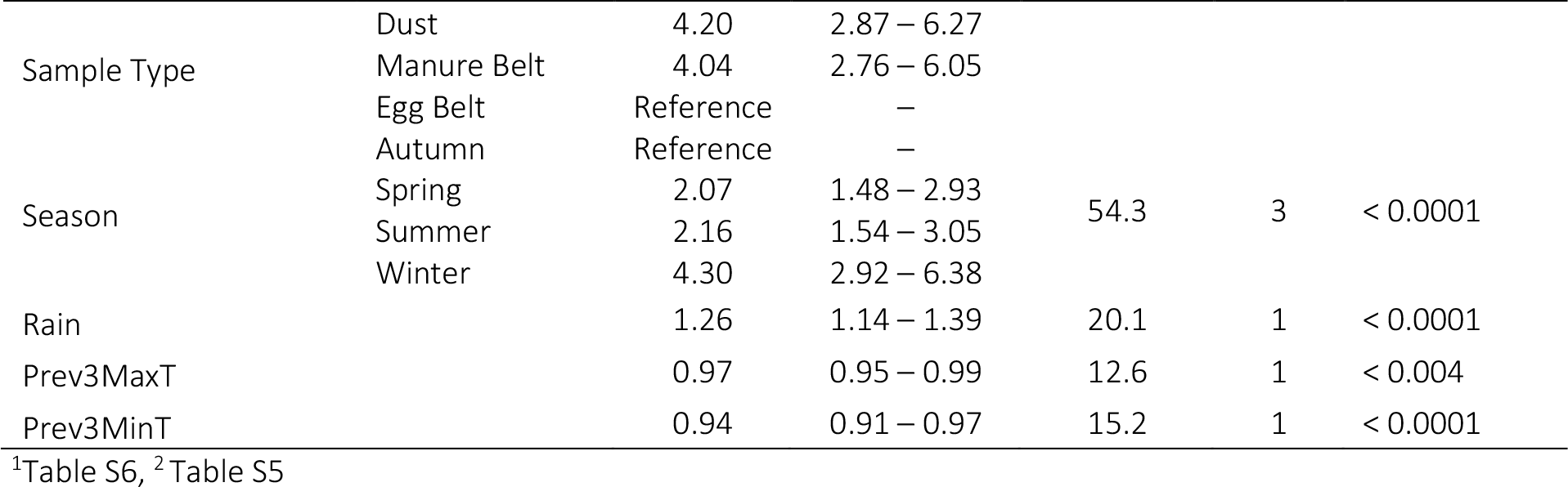
Univariate results for colony cage sheds - all statistically significant variables

**TABLE 8.**
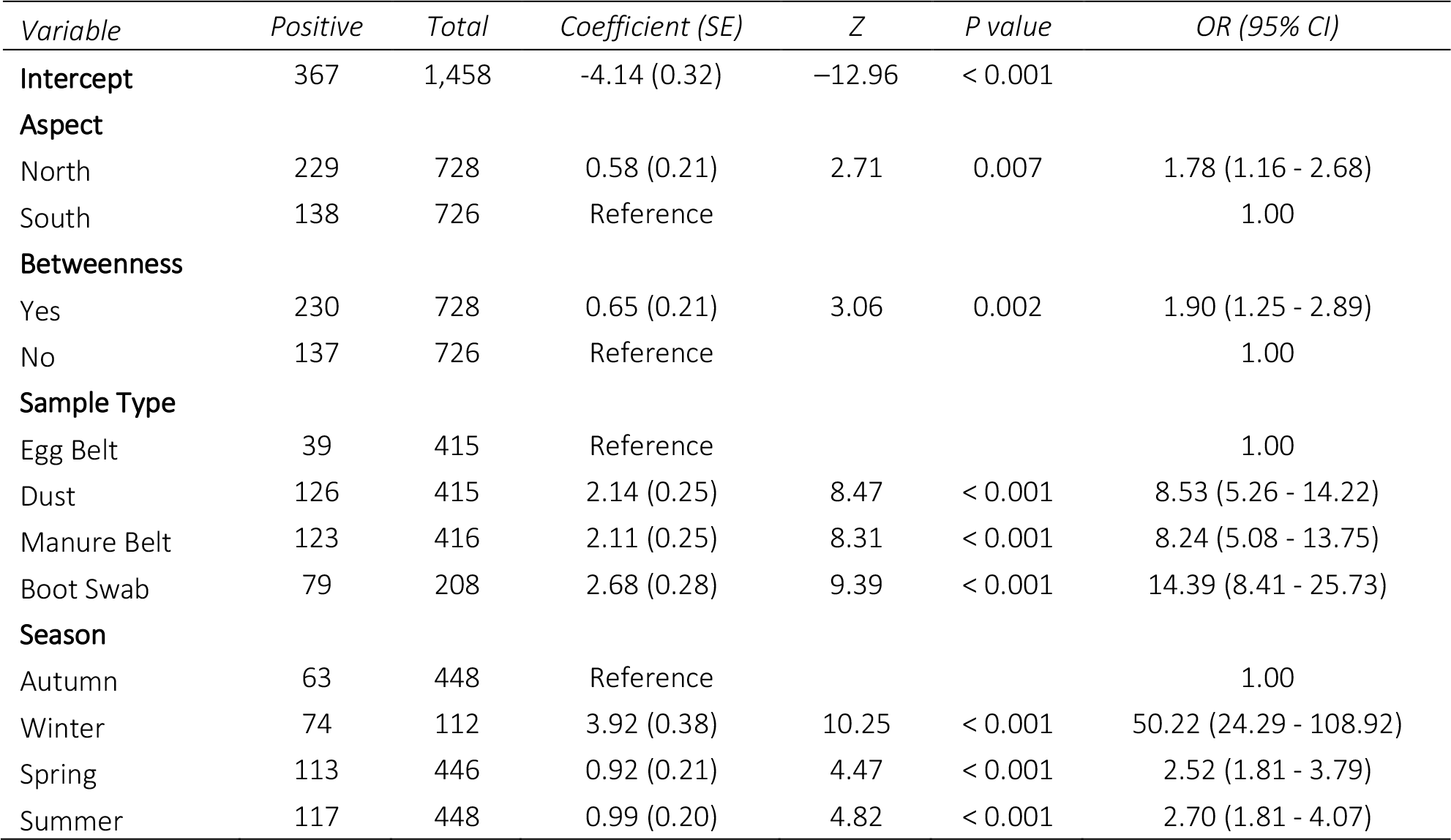
Estimated regression coefficients and standard errors for the final mixed-effects logistic regression model of environmental risk factors to predict the number of *S. enterica* positive environmental samples from a longitudinal investigation of caged chicken flocks (2013 - 2018).

Two-level hierarchical models with random effects considered to account for both sample event– and shed– level effects were built. The final most parsimonious model accounted for the difference in the shed-level effects as a fixed term, as there was no significant difference between the two models when both levels were included as random effects (Log-Lik (10) = −98.2, P = 1.0). The Hosmer-Lemeshow goodness of fit test was significant at small values of g = 5 to 10 (*χ*^2^ (3) = 2.18, P = 0.54, and *χ*^2^ (8) = 13.85, P = 0.086) indicating a relatively good fit to the data with this model. The residual variation due to unknown shed effects (variation partition coefficient) was estimated as (1.858/1.858 + 3.29) = 0.39.

The sensitivity and specificity of the model was estimated as 0.87 and 0.77 respectively. The area under the ROC curve or diagnostic accuracy (FIG S2) was estimated as 0.88 (95% CI [0.86, 0.90]), indicating a good prediction of the outcome by the model. Samples that were collected from a location on the north side of a shed (P <0.007) or between a shed (P < 0.002) were more likely to be positive for *S. enterica* serovars.

Both the location of sample collection and sample type were important but only sample type was included in the final model. The odds of a sample being positive was greater for boot swabs (P < 0.001) than any other sample type. As each sample type was collected by location it could be interpreted as a proxy variable for location. Individual weather variables were not significant in the final model (TABLE S9) but the effect of season was very important as samples collected in the winter months (P <0.001) were more likely to be positive than those collected in other season.

## DISCUSSION

This study investigated the performance of environmental sampling for the detection of salmonellae in caged environments under Australian environmental conditions. Sample size estimations demonstrated that at least 28 samples were required to detect salmonellae at a design prevalence of 1% with an imperfect test. As individual bird sampling for salmonellae is known to be less sensitive than environmental sampling (4), the number of birds to sample was estimated using the strictest criteria, proof of freedom of disease. This comparison (bird vs environment) was made to demonstrate the relative efficiency of environmental sampling versus individual bird sampling as a screening tool.

Both the logistics and practicality of selecting and testing hundreds of birds from each flock at regular intervals throughout production precludes this from being a real option for routine surveillance purposes. It also highlights that the number of samples (birds) required at very low flock prevalence is greater than the design criteria when using an imperfect test. Under these environmental conditions it is impossible to sample sufficient birds to determine the lowest flock prevalence of 1 positive unit per 100 units. This has important implications for confirmatory testing of the *S. enterica* status in flocks after a positive environmental sampling result, particularly when the flock prevalence is low.

Analysis of the field results demonstrated that, not only is the number of samples taken critical for detecting salmonellae within a caged environment, but where the samples are collected is also important. Samples were more likely to be positive in the winter and univariate analysis indicated decreasing temperature and increasing moisture also increased the odds of detection. Critically in the seasons studied, the winter was dry and cool (2.00 mm, 7.9-21.9°C) but not wet. These results are consistent with other studies that have demonstrated a seasonal effect on the detection of *Salmonella* spp. (9).

Despite the small number of sheds intensively sampled, many samples were collected frequently from the same locations within the sheds. The analyses demonstrated that salmonellae were heterogeneously distributed within the shed and that the distribution differed by serovar. The absence of spatial autocorrelation by location supports the conclusion that specific micro-environmental conditions are likely to be important in the survival or persistence of *S. enterica* within the shed. The presence of different serovars in specific locations and sample types supports the hypothesis that there are differences between bird-contact and other environmental surfaces and the detection of *S. enterica*.

Consequently, it is critical where sampling is conducted, because the presence or survival of salmonellae within the shed does not appear to be a random event. The effectiveness of detecting salmonellae varied by shed and this variation was related to the overall sample prevalence in the flock and therefore level of contamination in a particular shed. As the shed environment accounted for 60% of the variation in the sample prevalence within the shed, it is critical to consider the shed environment and its structure (bird contact versus non-contact surfaces) when designing sampling methodology.

The use of a multi-sample type sampling strategy is not novel and this approach was used to allow comparison with international studies using similar methods. However, the strategy for collecting samples was novel. A critical gap identified in the literature was where to collect samples from (other than randomly) and there is robust discussion about whether it is better to take a small sample more frequently from multiple locations rather than a large pooled sample of, for example, dust (1, 5).

Samples were deliberately collected systematically from within the shed at person height (tier two of each frame) to ensure that all frames of birds were sampled within easy reach, assuming dust samples are heavy and that any dust present at one level is from both the tier sampled and possibly the tiers above. Birds at this height are exposed to more interaction with people as they are in easy view and it may be hypothesised that they experience more stress, resulting in more shedding of salmonellae.

Multiple sites, and thus different sample types, were selected from each frame to maximize the opportunity to detect salmonellae. Manure belt samples were obtained at the end of each frame. Samples were obtained from egg belts from one tier of each frame and dust was collected from the same tier. Floor samples were collected from each half of the shed. It was hypothesized that these sites, except boot swabs, were less likely to be cross contaminated by people movements in and out of the shed.

We included in this surveillance design the use of boot swabs on concrete floors. Boot swabs are typically used on litter or slatted floors in free range layer sheds or broiler production sheds (14) but this study demonstrated that they are also a very sensitive method of sampling in the cage environments. Boot swabs were more likely to test positive for *S.* Infantis than for *S.* Typhimurium, which may reflect better survival of *S*. Infantis in the floor environment. Finally, manure samples were collected by directly sampling the end of the manure belts, where they are more easily accessed, rather than collecting a pooled sample of faeces. Depending on the shed design, access to fresh manure under cages can be very difficult, additionally the timing of sampling is critical as manure belts may have been cleaned just prior to sampling, which will limit access to an adequately representative manure sample. Manure belt samples were as effective as dust samples for salmonellae detection and there was no difference in the detection of the two *S. enterica* serovars. Dust or manure belt swabs were easier to collect and preferable to handling large quantities of manure for collection, laboratory testing and disposal.

These specific sampling sites were chosen to ensure that the location and type of sampling was repeatable on all sampling occasions by all samplers. It is important to note that, while obvious, birds housed in caged environments cannot roam freely within the whole shed space. Additionally, birds are housed both vertically and horizontally. This means that if infection within the flock is not homogeneously distributed then failing to sample the entire shed space may bias the sampling results, particularly if the prevalence is low. Other studies have reported that sample prevalence may be lower in birds housed in higher tiers of the frames (23).

The positive predictive value of the overall sampling strategy was high (98%), even when accounting for an imperfect test. Each of the sample types performed well with the key exception of egg belt swabs in this study, where the likelihood of detection was lower. Egg belts in these sheds were constructed of a polyethylene plastic material and this may have hindered detection, because the surface is prone to rapid drying and may not be a suitable environment for long term survival. Also, it is likely that insufficient egg belt surfaces were swabbed. The egg belt is the only surface in these sheds that was not as readily cross contaminated with dust or environmental material as the other exposed surfaces because it is protected from above.

A number of other potential factors were deliberately not included in this study but may affect the presence of *S. enterica* within the shed as a consequence, or reflection of, bird stress. The higher prevalence of positive environmental samples on northern or sheltered aspects (hotter in southern hemisphere) of the shed may be an indicator of increased stress in the flock, with consequential increased shedding in that part of the flock, or it may indicate preferential long term survival of salmonellae under those environmental conditions. Unfortunately, internal shed environmental records were only available for the whole shed, rather than for specific areas of the shed, so these effects could not be investigated further.

These findings are particularly relevant to other housing designs, such as aviary housing where birds may be housed freely but have both vertical and horizontal access within a shed environment, in a similar organisational arrangement to birds in caged sheds, which makes sampling individual birds significantly more challenging. The sampling principles as those described here are easily applicable to these environments with the exact same sampling considerations applied to the space.

## CONCLUSION

This study confirmed that the number of samples chosen is important for *S. enterica* detection, but of greater importance is how and where samples are collected from within a caged flock shed environment. *S. enterica* distribution within a shed is not homogenous and the location of *S. enterica* serovars within a shed does not appear to be random. Factors influencing detection include the season, the weather (low rainfall and moderate temperatures) and the shed aspect (north or south side) or sheltering by another building. A multi-sample type approach has high positive predictive value even when the diagnostic test sensitivity is low and the environmental prevalence is low.

*S. enterica* serovars were detected in different spatial locations within the shed indicating that specific micro-environments may enhance survival, and possibly multiplication, and subsequent detection. All these factors should be taken into account, when designing a surveillance strategy for the detection of salmonellae in caged flocks. In sheds of this type, multiple samples should be collected from different areas, preferable in close contact with the birds, focusing on manure belts, dust and boot swabs. If resources limit the number and type of sample that can be collected and processed, then boot swabs are a good first choice of sampling material. Regardless of the sample type chosen, sampling must be conducted to ensure that the whole shed space is sampled as homogeneously as possible.

## MATERIALS AND METHODS

### Study population

Nineteen cage layer production flocks from farms in Victoria, Australia, with a history of infection with salmonellae were purposefully selected for sampling. Each flock was housed as a single age group, with all-in all-out management. Each flock was placed in the shed after cleaning and disinfection of the shed and prior to the onset of lay at 16 - 19 weeks of age.

Samples were collected over a 4 year period 2014 and 2018, with the following months aggregated for each southern hemisphere season: Summer, December – February; Autumn, March – May; Winter, June – August; and Spring, September - November. This study was conducted under normal farming operating conditions to evaluate routinely conducted sampling procedures for surveillance purposes. All environmental sampling was conducted as part of routine production standard operating procedures in accordance with standard industry procedures and guidelines (15, 24–26). As all samples were collected as part of routine veterinary care and agricultural practice this study did not require ethics approval (27).

### Bird housing and shed design

Birds were housed in compliance with the Australian model code of practice for poultry (25) and state legislation (28). Birds were housed in multi-tiered frames in either enriched colony cages with one cage width per frame, or conventional cages, two cages back to back per frame width. Colony caged sheds housed ∼24,000 birds in eight frames, 3 – 4 tiers high, in 48 colonies each containing ∼500 birds. Conventional cage sheds housed ∼65,000 birds in 5 frames 6 tiers high with each cage containing ∼5 - 6 birds. Cages were of European design and construction from two major suppliers of cage equipment.

Sheds of the same type were identical in design and construction, only varying in their positioning next to another shed and whether the long length of the shed was exposed or sheltered by another shed, FIG. 1. Sheds were ∼130 m × 30 m in dimension with concrete floors. Walls had short concrete walls from the floor (∼ 600 mm) topped by an aluminium bonded insulated panel to the ceiling with no exposed joists or framing. Ceilings were constructed of the same material as the walls and greater than 6 m in height. Sheds were climate controlled with evaporative cooling (cool pads and fans) and heat exchange for warming. Eggs were automatically collected from nest boxes or cages to an egg belt delivered to a vertical elevator at the end of each frame where they were travelled to a central egg packing room. Manure was collected on belts under the cages and removed at least weekly from one end of the shed. All sheds were placed with the same orientation to the sun, with the long length of the shed positioned west-east (Southern Hemisphere).

**FIG. 1.**
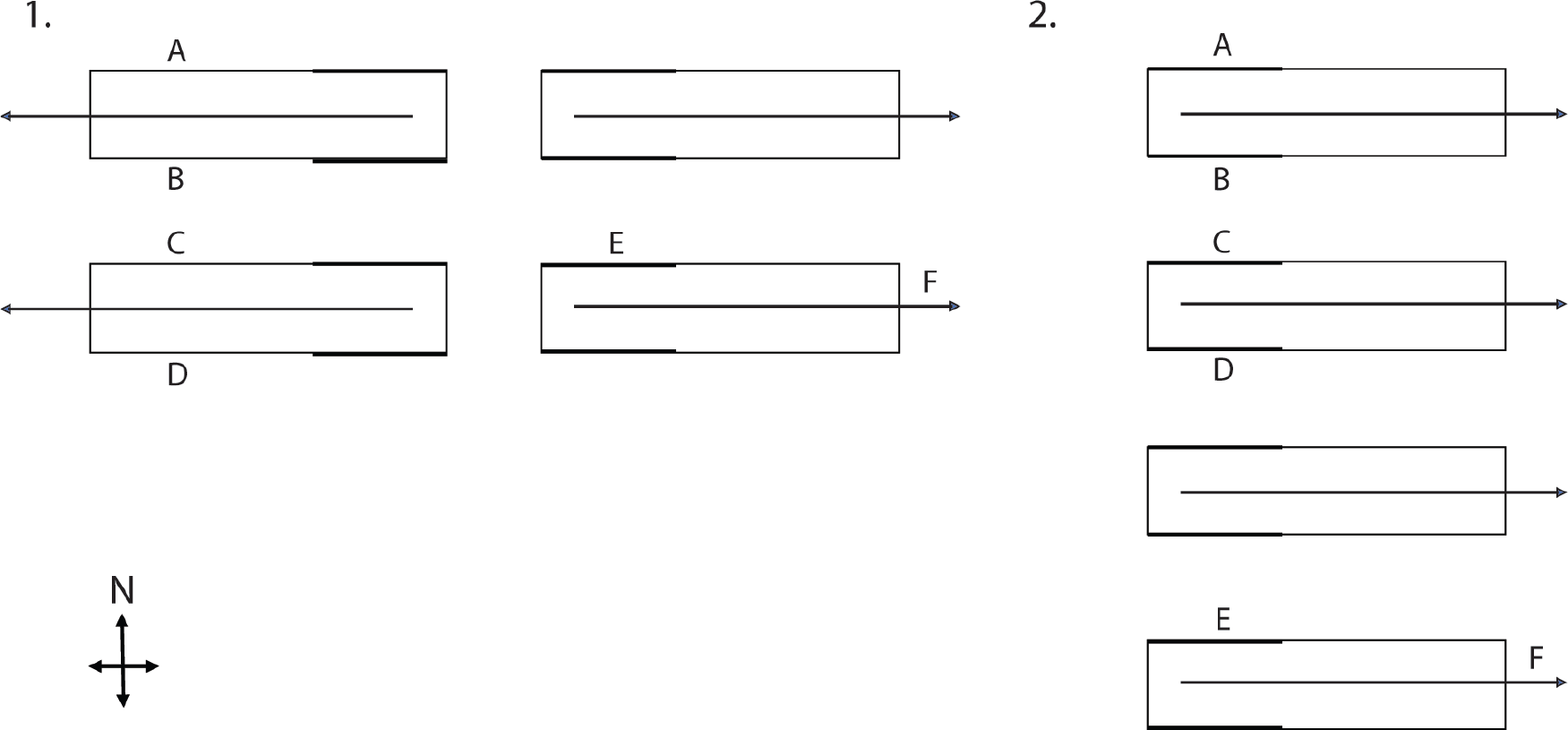
Shed arrangement and orientation. (1) Side by side shed arrangement, (2) Linear shed arrangement. A. Northern aspect, B. Southern aspect between shed, C. Northern aspect between shed, D. Southern aspect, E. Cool pad location, F. Tunnel ventilation direction of airflow.

### Sample size and prevalence calculations

Sample size was estimated for three design prevalence estimates of 1, 5 or 10 *S. enterica* positive units per 100 units at risk (1. Birds, 2. Cages or 3. Environmental sites) with 95% confidence that the estimated prevalence was within 5% of the true prevalence in the unit tested, assuming an imperfect test.

Two sample size calculation methodologies were used; to estimate the number of birds to sample, freedom from infection using an imperfect test was used (29, 30), and for the number of cage or environmental sites to be sampled the true prevalence assuming an imperfect test was used (21, 31). For sample size calculations, diagnostic test sensitivity and specificity for *S. enterica* by culture was assumed to vary between 0.88 - 0.98 and 0.99 - 1.00 respectively (32–34).

Prevalence estimates based on the results of testing (apparent prevalence) were distinguished from prevalence estimates corrected on the basis of imperfect diagnostic test performance (true prevalence) (35, 36). For the calculation of true prevalence, test sensitivity was assumed to be 0.88 and specificity 0.995.

### Environmental sample sites

A total of 168 potential sampling sites were identified within each shed. These sampling sites included manure belts, frame surfaces, feed lines or feeders, nest box surfaces, fan covers, floor and wall surfaces and egg belts. Sites were identified as either bird contact areas (hypothesized to reflect flock infection status); those immediately within the bird surrounds such as manure belt, egg belt, nest box surfaces or the framing, or, non-bird contact areas (hypothesized to reflect shed contamination status) indirectly contaminated by dust, dander or feed residue such as walls, floors, fan covers, feed lines. Potential bird contact sampling locations are indicated in FIG. 2.

**FIG. 2.**
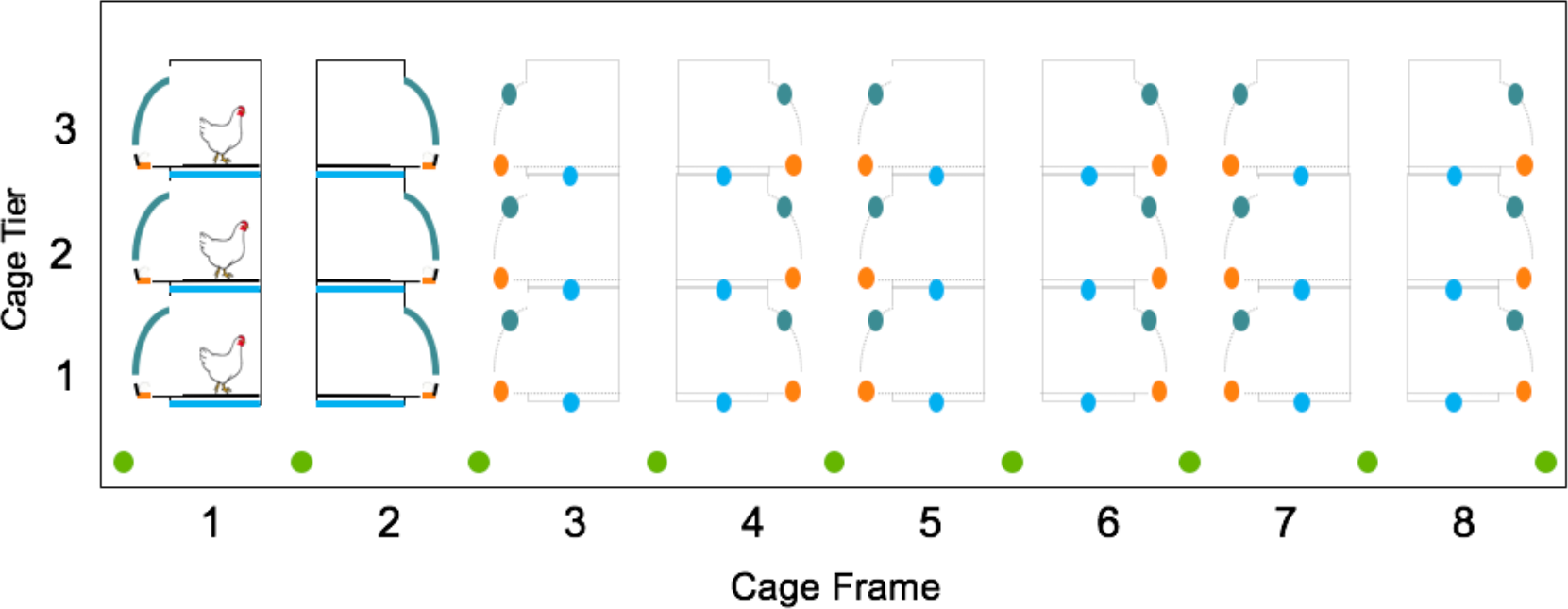
Bird contact sampling locations within cage house for each cage tier and cage frame (colony cage style illustrated). Boot Swab = Green, Manure Belt = Blue, Egg Belt = Orange and Dust = Dark Blue.

**FIG. 3.**
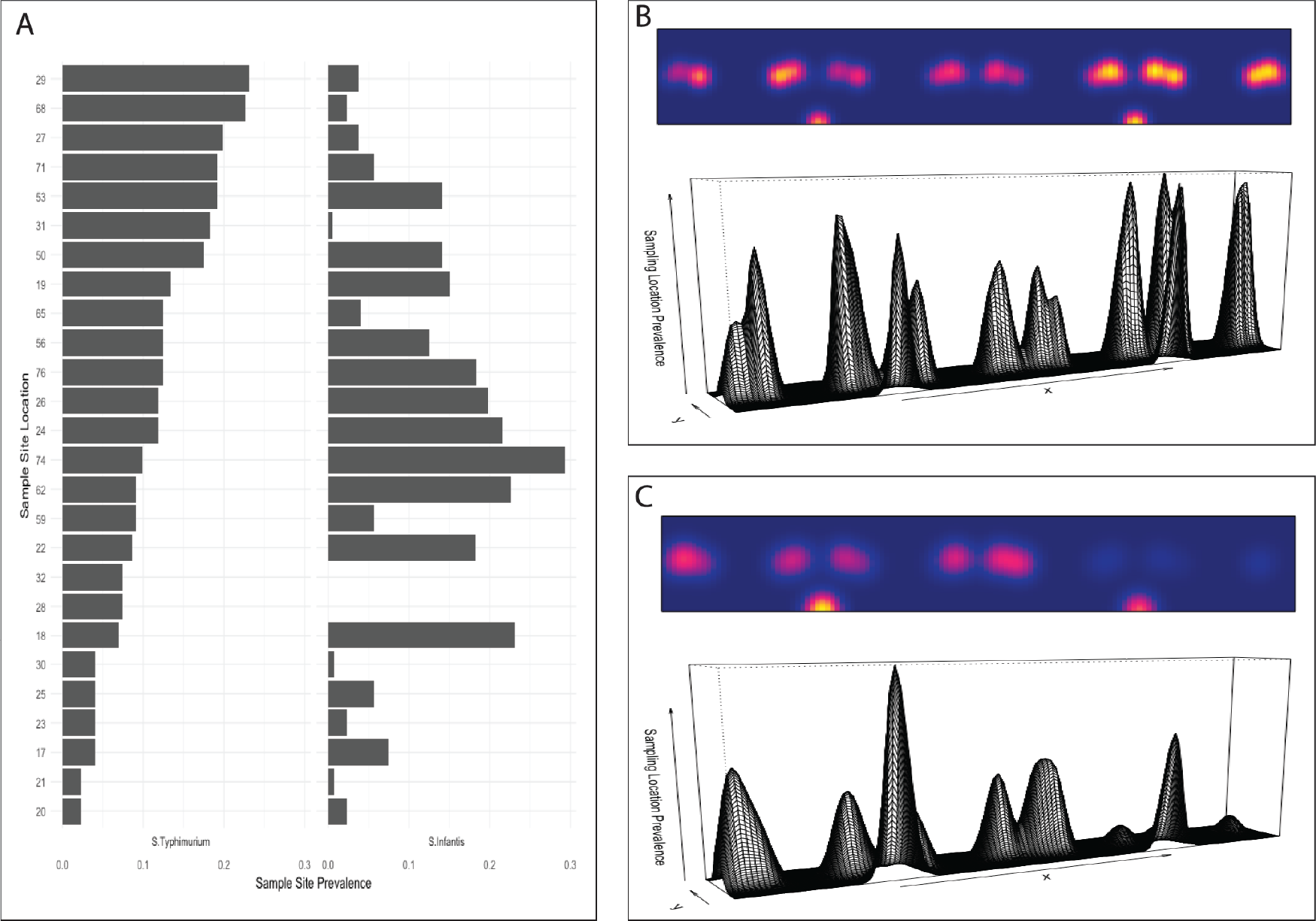
Heterogeneity of *S.* Typhimurium and *S*. Infantis distribution within a colony cage environment. The environmental prevalence for each serovar is calculated from all samples collected from that sampling location. (A) *S. enterica* prevalence at the sampling location within the shed environment for *S*. Typhimurium and *S.* Infantis separately. Length of the bar indicates serovar prevalence at that sampling location. Sampling location is ordered by the site with the highest *S.* Typhimurium prevalence. (B) Heat map indicating *S*. Typhimurium prevalence in a two-dimensional space at each sampling location. The warmer the colour the greater the prevalence at that location. Corresponding perspective plot of *S*. Typhimurium prevalence, height is proportional to the prevalence at the sampling location. (C) Heat map indicating *S*. Infantis prevalence in a two-dimensional space at each sampling location. The warmer the colour the greater the prevalence at that location. Corresponding perspective plot of *S.* Infantis prevalence, height is proportional to the prevalence at the sampling location.

### Environmental sampling methodology

Within each cage shed, 28 or 29 samples (8 – 10 egg-belt, 8 – 10 dust, 5 – 8 manure belt and 4 boot swabs) were collected on each sampling occasion. Four 10 x 10 cm cotton gauze swabs, pre-moistened with buffered peptone water, were used to collect each surface sample. Manure belt samples were collected by wiping the exposed edge of all belts, at one end of the shed, avoiding excessive shed “dust” and debris, concentrating on exposed faecal material on the leading surface and immediately under the belt rather than the top exposed surface. Egg belts were sampled by wiping the length of the bottom surface of the egg belt from a single tier. Dust samples were collected by wiping the surface of the nest box the length of the frame. Clean dry boots, with new plastic boot covers were worn on entry to the shed. Two pairs of boot swabs were worn while walking within the shed during sample collection. The second pair was exchanged during the mid-point of sampling the shed. Each sample-type was pooled separately by frame and cage row into a Whirlpak™ bag and identified by shed, flock, sample type and sample location. Samples were refrigerated immediately after collection.

### Sample processing and microbiology

All samples were collected and returned to the laboratory for processing the same day. Each sample was processed separately, no samples were pooled. All samples were cultured in accordance with the Australian Standard 5013.10 (ISO 6579: 2002 MOD) for the detection of *Salmonella* spp. from environmental samples (37). The full sample processing methodology is available online (38) and details are provided in the supplementary material (Microbiological methods).

### Statistical analysis

All analyses were conducted in the R statistical package unless otherwise stated (39). Spatial analysis was conducted using the following packages: “spsurvey” (40), “rgeos” (41), “sparr” (42) and “spatstat” (43, 44). Measures of association, odds ratios and positive predictive values were estimated using “epiR” (45). The R packages “nlme”, “lme4” and “aod” were used for hierarchical mixed effects model building (46–48).

### Assessment of sample site homogeneity

The spatial distribution of *S. enterica* serovars within each shed was evaluated by aggregating all sampling events for each shed. Spatial heterogeneity was evaluated by spatial point pattern analysis and calculating the spatial density (kernel) of each of the sampled sites using “spatstat” (43, 44). The distribution of *S. enterica* within the shed was visualized using heat maps to plot the calculated density at each sampling location within a two-dimensional representation of the shed environment. Spatial autocorrelation was evaluated using Moran’s I from the “ape” package (49) and the resulting variogram visualised using “geoR” (50).

### Univariate Analysis

There were two outcomes of interest: 1. Binary, the presence (1) or absence (0) of salmonellae in a sample; and 2. The true sample prevalence at a given sampling event. Only explanatory variables associated with the sampling methodology and the housing environment were considered in this analysis. A full description of each variable considered for analysis is available in Table S9.

Measures of association between the binary outcome of interest and the explanatory variables were computed using the odds ratio. For categorical or continuous variables, simple linear or logistic regression was used where appropriate. If an explanatory variable was statistically associated with the binary outcome of interest in the univariate analysis (P < 0.10) it was considered for inclusion in the multivariable analysis.

### Multivariable hierarchical mixed-effects logistic regression

Two fixed-effects multivariable logistic regression models were built where the probability of a sample being positive for *S. enterica* at a sampling event was parameterised as a function of the significant explanatory variables (P < 0.10) identified in the univariate analysis. Model 1 was built using all available data for all 19 flocks. Model 2 was built for the subset of 4 flocks that had intensive sampling for the life of the flock; the first 13 sampling events of these four flocks were included in this analysis.

The models were built in a stepwise manner with all variables included initially and non-significant explanatory variables removed sequentially from the model, until all variables retained in the model were significant at *α* < 0.05. Variables removed from the model were retested in the final model and retained if their inclusion changed the regression coefficients significantly, using the Wald test. Where no significant difference was observed between two models then the most parsimonious model was selected. Biologically plausible interactions were considered. Autocorrelated measures were identified and tested individually in the model, only one was kept in the model if it remained significant.

Due to the hierarchical and longitudinal nature of the data (TABLE S10) samples clustered within sampling events, and sampling events clustered within sheds and sheds within farms, the model was extended to include sample event–, shed– and farm–level random effect terms, where appropriate. Factors influencing the detection of *S. enterica* were categorized into three classes: those operating at the sample level, the sample event level and the shed level. Sample level effects considered included the sample type, sampling location, whether the sample was collected from the northern aspect of a shed or whether the sampling location in a shed was between two sheds. Sampling event level effects included the month of sampling, the season of sampling event and the weather (including rainfall, temperature and solar radiation at the time of sampling or the three weeks prior to sampling). For this analysis, the shed level effects considered were the Salmonella status of the previous flock and the presence of salmonellae in the shed prior to the onset of the study and the flock.

The assumptions of normality and homogeneity of variance of the final model were checked using frequency histograms of the residuals, and plots of the residuals versus predicted values. The Hosmer-Lemeshow goodness of fit test was used to evaluate the fit of the final model and the diagnostic accuracy was computed by evaluating the ROC curve from the model estimates.

## ACKNOWLEDGEMENTS

This work was supported by grants from the Cybec Foundation, Victoria, Australia. We are extremely grateful to the poultry producers that allowed us access to their properties for this research and their willingness to participate in this study.

## SUPPLEMENTARY MATERIAL

Supplementary material for this article may be found in Supplementary File 1

